# Collagen-binding integrin α11β1 contributes to joint destruction in arthritic hTNFtg mice

**DOI:** 10.1101/2022.01.14.476301

**Authors:** Adrian Deichsel, Anna De Giuseppe, Isabel Zeinert, Kerstin Rauwolf, Ning Lu, Denise Beckmann, Annika Krause, Beate Eckes, Uwe Hansen, Daniel Kronenberg, Donald Gullberg, Thomas Pap, Adelheid Korb-Pap

## Abstract

**Background:** In rheumatoid arthritis (RA), fibroblast like synoviocytes (FLS) undergo a “tumor-like” transformation, wherein they develop an aggressive phenotype that is characterized by increased adhesion to components of cartilage extracellular matrix (ECM) and that contributes extensively to joint destruction. The collagen binding integrin α11β1 was previously shown to be involved in similar processes in cancer-associated fibroblasts mediating tumorigenicity and metastasis in certain tumors. Therefore, this study aimed to study the role of integrin α11β1 in RA and to characterize the effects of α11β1 deficiency on the disease course and severity in arthritic hTNFtg mice.

**Methods:** The expression levels of integrin α11β1 were analyzed by immunohistochemistry, immunofluorescence, and western blot analysis in synovial samples and FLS of patients with RA and osteoarthritis (OA) as well as in samples from wild type (wt) and arthritic hTNFtg mice. Furthermore, the subcellular expression of integrin α11β1 was investigated in co-culture experiments with cartilage explants and analyzed by transmission electron microscopy. To investigate the effects of integrin α11β1 deficiency, *itga11*^*-/-*^ mice were interbred with hTNFtg mice and disease severity was assessed by clinical scoring of grip strength and paw swelling over the disease course. Hind paws of 12-weeks-old mice of all genotypes were analyzed by µCT imaging followed by stainings of paraffin-embedded tissue sections with Toluidine-blue and tartrate-resistant acid phosphatase (TRAP) to evaluate established parameters of joint destruction such as inflammation area, cartilage destaining, FLS attachment to the cartilage surface, and bone damage.

**Results:** Expression levels of integrin α11β1 were clearly elevated in synovial tissues and FLS from RA patients and hTNFtg mice, compared to the controls derived from OA patients and wt mice. Interestingly, this expression was shown to be particularly localized in focal adhesions of the FLS. As revealed by transmission electron microscopy, integrin α11β1 expression was particularly evident in areas of direct cellular contact with the ECM of cartilage. Evaluations of clinical scorings and histomorphological analyses demonstrated that *itga11*^*-/-*^hTNFtg displayed alleviated clinical symptoms, higher bone volume, less cartilage destruction and reduced FLS attachment to the cartilage in comparison to hTNFtg mice.

**Conclusions:** The collagen-binding integrin α11β1 is upregulated in the context of RA and its deficiency in mice with an inflammatory hTNFtg background leads to a significant reduction in the arthritic phenotype which makes integrin α11β1 an interesting target for therapeutical intervention.

## Background

Rheumatoid arthritis (RA) is a chronic systemic inflammatory disease with a high prevalence and socioeconomic burden affecting joints in a characteristic symmetrical pattern. If untreated, RA leads to massive irreversible destruction of articular structures such as cartilage and bone. It has been shown that, in addition to the influx of immune and inflammatory cells, fibroblast-like synoviocytes (FLS) can be attributed a key role in the pathogenesis of RA, as these cells are known to contribute significantly to the development, progression and chronic course of this disease [1]. A major feature of FLS in RA is their stable activation and transformation into an autonomously aggressive phenotype. This, among other features, leads to an increased expression of adhesion molecules, triggering further destructions of cartilage as these facilitate the binding of RA-FLS to components of the extracellular matrix (ECM) [2,3].

In this context, different matrix adhesion molecules such as integrins and syndecans were shown to play important roles in the attachment of RA-FLS to collagens [4-6]. One member of the β1-family of integrins that has not been investigated in detail in RA, is integrin α11β1. Integrin α11β1 is encoded by the gene *itga11* and one of four known integrins (besides α1β1, α2β1 and α10β1) to bind to collagen [7, 8]. First discovered in 1995 by Gullberg et al, it was shown to be expressed mainly on cells of mesenchymal origin, notably fibroblasts [9-11]. Integrin α11β1 was demonstrated to mediate not only cellular adhesion but also cell migration, collagen reorganization and the expression of matrix modifying enzymes [7, 12, 13]. In the tumor-context, integrin α11β1 as expressed in cancer-associated fibroblasts (CAFs) within the tumor stroma has been shown to positively influence tumor growth, metastatic potential and negatively affect disease outcome [14-16].

Due to the similarities between tumor cells and RA-FLS, which amongst others are characterized by increased adhesion, migration and invasion capacities, we hypothesized that α11β1 not only plays a role on CAFs in different cancer types, but could also trigger disease severity also in RA.

## Methods

### Human synovial tissues and fibroblast-like synoviocytes

The ethics committees of the Medical University of the University Hospital Muenster approved all studies with human samples. Samples of synovial tissues from subjects with RA or OA (according to the 1987 revised American College of Rheumatology criteria for RA and OA [17]) were obtained as operational waste at joint replacement surgery and all subjects gave informed consent prior to surgery.

RA-FLS and OA-FLS were isolated by enzymatic digestion using the collagenase type IV (Worthington Biochemicals) and cultured in 10% heat-inactivated FCS-supplemented Dulbecco’s modified Eagle’s medium at 37°C and 5% CO_2_. Cell suspension was centrifuged at 1500 rpm and RT for 5 minutes, the pellet was resuspended with DMEM and FLS were cultured under standard conditions. To eliminate initial contaminations with other cells, only cells at passages 3 to 5 were used for experiments.

### Animals

The hTNFtg mice carrying the transgene from human tumor necrosis factor-α (strain Tg197; C57BL/6 genetic background; obtained from Alexander Fleming Biomedical Science Research Center, Vari, Greece) and α11β1 mice (kindly provided by D. Gullberg, Bergen, Norway) were described previously [18, 19]. Both mouse strains were interbred within the C57BL/6 genetic background. The genotype was confirmed by polymerase chain reaction (primer sequence, see Tab. 1). Mice were scored on a weekly basis up to an age of 12 weeks to evaluate arthritis symptoms. The evaluation was based on a scoring range from 0 (no symptoms) to 3 (severe symptoms), including grip strength, paw swelling and weight [20]. All animal procedures were approved by the State Office for Nature, Environment and Consumer Affairs (Landesamt für Natur, Umwelt und Verbraucherschutz LANUV), Germany (reference numbers AZ 84-02.04.2015.A511).

### Isolating and fibroblast-like synoviocytes from mice

Mice were sacrificed in accordance with the German animal welfare act using carbon dioxide (CO_2_). Skin and nails of the hind paws were removed and the larger ligaments were dissected. Finally, the hind paws were dislocated and paws were digested with 1mg/ml collagenase (Collagenase Type IV, Worthington Biochemicals) in Dulbecco’s modified Eagle’s medium (DMEM) for 1 h at 37°C. After digestion the cell suspension was centrifuged at 1500 rpm and RT for 5 minutes. The supernatant was discarded, the pellet resuspended in DMEM supplemented with 10% heat-inactivated fetal calf serum (h-FCS) and 1% Penicillin-Streptomycin. Isolated fibroblasts were cultured at 37°C and 5% CO_2_, experiments were performed between passage 3 and 5.

### Cartilage attachment assay and transmission electron microscopy

As previously described [4], freshly isolated cartilage of the femoral head of 4-6 weeks old mice and isolated wt and hTNFtg FLS were co-cultivated in FLS medium for three days followed by transmission electron microscopy and immunogold-mediated detection of integrin α11β1.

### Preparation of human and murine tissues for histology

Human samples from RA and OA patients as well as the hind paws from twelve weeks old mice were fixed in 4% paraformaldehyde overnight at 4°C and decalcified in 20% Na-ETDA (*AppliChem*) for eight weeks. Afterwards, decalcified tissues were dehydrated and embedded in paraffin. Paraffin-embedded human tissues and hind paws were cut into 5 μm sections with the Microtome HM355S (*Thermo Fisher Scientific*) and transferred onto microscope slides.

### Immunohistochemistry staining of human and murine synovial tissue

Sections of decalcified, paraffin-embedded hind paws and human synovial tissues were deparaffinate in xylene and rehydrated in decreasing concentrations of ethanol. Subsequently, sections were incubated in distilled water and washed in PBS. Peroxidase activity was blocked with a 30% hydrogen peroxide solution in methanol. The sections were pretreated with 1x trypsin for 10 min at 37°C and blocked with 20% normal horse serum for 1 hour. Human tissues were stained with a sheep polyclonal antibody to integrin α11 (*R&D Systems*) and murine tissues with a rabbit polyclonal antibody to mouse α11 generated and kindly provided by Donald Gullberg (Department of Biomedicine, University of Bergen, Norway). As secondary antibody, a biotinylated anti-sheep IgG or anti-rabbit IgG (Vector Laboratories) were used. The stainings were performed using the Vectastain ABC peroxidase kit and DAB substrate kit (*Vector Laboratories*). Counterstaining was conducted with Mayer’s Haemalaun (*Sigma-Aldrich*). Sections were mounted with Dibutylphtalate polystyrene xylene (DPX) for microscopy.

### Toluidine-blue and haematoxylin-eosin (HE) staining of paraffin sections

Paraffin sections were deparaffinate in xylene and rehydrated in decreasing concentrations of ethanol. Subsequently sections were incubated in distilled water and stained with Toluidine-blue (*Sigma-Aldrich*) or Mayer’s Haemalaun (*Sigma-Aldrich*) and eosin Y (*Sigma-Aldrich*). Stained sections were dehydrated in increasing concentration of ethanol and incubated in xylene followed by mounting the slides with DPX.

### TRAP staining

The tartrate-resistant acid phosphatase (TRAP) kit (*Sigma-Aldrich*) was used for osteoclasts detection on paraffin sections of twelve weeks old hind paws using the TRAP kit following the manufacturer’s instructions.

### Integrin α11β1 expression levels in murine and human FLS

FLS were lysed in radioimmunoprecipitation assay (RIPA) buffer. Protein concentrations were determined by a bichinonic acid (BCA) protein assay kit (*Thermo Fisher*) according to manufactures instructions. The protein extracts were resolved by a dodecyl sulfate polyacrylamide gel electrophoresis using a 12% separation gel. Gels were transferred to a polyvinylidene difluoride (PVDF) membrane (*GE Healthcare*) in a Trans-Blot Turbo device (Bio-Rad). Integrin α11 expression levels were detected by rabbit polyclonal antibodies to human or mouse integrin α11 (provided by Donald Gullberg, Department of Biomedicine, University of Bergen, Norway) and the polyclonal anti-rabbit immunoglobulins/HRP (*Dako*). Images were analyzed using the gel analyzing tool by *ImageJ, version 2*.*1*.*0/1*.*53c*.

### Immunofluorescence staining of human and mouse FLS

Cells were seeded on sterile glass coverslips coated with bovine collagen coating solution (*Cell Applications, INC*.) for improved attachment. FLS were incubated in DMEM supplemented with 10% h-FCS at 37°C and 5% CO_2_ overnight. Thereafter, cells were washed in PBS and fixed for 20 minutes in 4% paraformaldehyde. Next, ammonium chloride was used to reduce the auto-fluorescence of the cells followed by permeabilization with 0.1% Triton X-100. Afterwards, FLS were incubated with 10% normal horse serum for 20 minutes at RT. Murine and human cells were stained with primary antibodies (polyclonal antibody for mouse and human α11; Donald Gullberg) for one hour and with the secondary Alexa Four 488 antibody (*Life Technologies*) for 30 minutes at RT. The cytoskeleton was stained with rhodamine phalloidin (*Invitrogen*) and nuclei stained with 4’,6-diamidino-2-phenylindole (DAPI) (*Invitrogen*). Mowiol was used as mounting medium.

### Histomorphometric analysis

Toluidine-blue stained sections were used for analyzing synovial inflammation, total cartilage area, cartilage damage, and attachment of FLS to the cartilage surface. Destained cartilage as a result of proteoglycan loss and cartilage degradation was quantified to the total amount of cartilage and indicated as a percentage. Synovial inflammation area was evaluated by relating the pannus tissue to the total tissue area expressed as a percentage, furthermore the length of the FLS attachment to the cartilage surface was evaluated. pannus tissue invading the cartilage. Quantification of the images were performed by using Zen Pro Software (*Zeiss*).

### Micro-computed tomographic analysis

The right hind paws from twelve weeks old mice were dissected from the leg, the skin and claws were removed and fixed overnight in 4% paraformaldehyde at 4°C. Hind paws were transferred in PBS and scanned with the *SkyScan 1176* (*Bruker, version 11*.*0*.*0*.*2*) at 40 kV tube voltage, 0.6 mA using an aluminium filter (0.2 mm thick) and 0.5° rotation steps. Associated software was used for the reconstructions (*NRecon, version 1*.*7*.*5*.*9*), 3-dimensional vizualisation (*CTVox, version 3*.*3*.*0r1412*) and analysis (*Data Viewer, version 1*.*5*.*6*.*6* and *CTAn, version 1*.*20*.*3*.*0*). BV/TV was measured from the tarsal bones 2-4 by drawing manually the borders of the bones and calculating the average.

### Statistical analysis

Graphs are represented as box-plots showing all data points with whiskers from minimum to maximum. The software *GraphPad Prism 9* was used for statistical analysis. Comparison of the different groups were performed by the two-tailed Mann-Whitney U test. P-values of less than 0.05 were considered to be statistically significant.

## Results

### Integrin α11β1 is upregulated in RA-FLS in the context of RA and in the hTNFtg mouse model

To assess the expression levels of integrin α11β1, paraffin-embedded human synovial tissues obtained from RA patients and - as a control - from OA patients undergoing joint replacement surgery were stained by immunohistochemistry for integrin α11β1. Tissue samples from RA patients displayed the characteristic formation of an invasive hypercellular pannus and a highly upregulated expression of integrin α11β1. In comparison, no pannus formation and very weak integrin α11β1 expression were observed in tissues from OA patients (Figure 1a). As a next step,we tested whether a staining pattern comparable to the human situation could be seen in an animal model of RA. To this end, decalcified, paraffin-embedded and sectioned hind paws derived from wild type (wt) and hTNFtg mice at an age of twelve weeks were used for immunohistochemistry stainings using a specific antibody against integrin α11β1. The hTNFtg mouse model is a well described mouse model for inflammatory polyarthritis and is characterized by a spontaneous joint inflammation due to genetic alteration leading to an ubiquitous overexpression of human TNFα [18]. Comparable to the staining pattern in human RA synovial tissues, hTNFtg mice also showed a strong upregulation of integrin α11β1 expression whereas only a faint staining for integrin α11β1 was found in wt section (Figure 1a).

**Figure 1.**
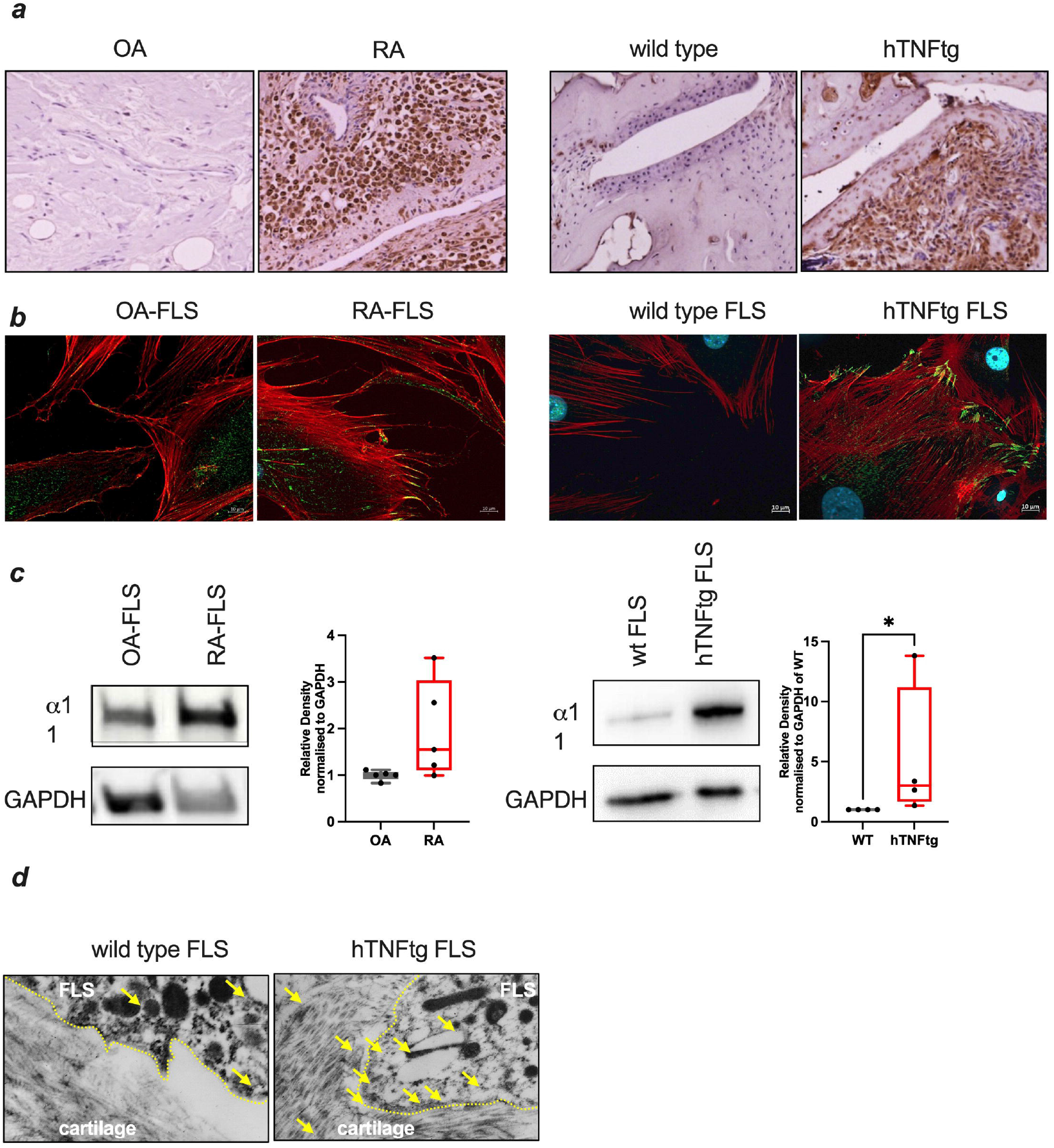
Expression of integrin α11β1 in inflammatory arthritis. Left side: Human tissue of OA and RA patients (n =5). Right side: Murine tissue of wt and hTNFtg mice (n =4). **a)** Immunohistochemistry stainings using specific antibodies against integrin α11β1 visualized by DAB. **b)** Immunofluorescence of FLS using specific antibodies against integrin α11β1 (green), F-Actin was visualized by rhodamine phalloidin (red) and nuclei by DAPI (blue) (error bars = 10 µm). **c)** Western blot analysis of integrin grin using specific antibodies against integrin α11β1 expression in FLS, GAPDH served as loading control. **d)** Immunogold-labelling of integrin α11β1 in wt and hTNFtg FLS co-cultivated with cartilage explants derived from murine femoral head.

As it is known that integrin α11β1 is mainly restricted to mesenchymal cells, FLS from RA and OA patients and from wt and hTNFtg mice were analyzed for their expression levels of integrin α11β1 as well as its subcellular expression pattern. In line with the previous *in vivo* data, RA-FLS and hTNFtg FLS showed an increased expression of integrin α11β1 compared to the controls, and integrin α11β1 was found primarily at focal adhesion sites (Figure 1b). Since both the immunohistochemical images of the paws and the immunofluorescence of the FLS showed an upregulation of integrin α11β1 expression under inflammatory conditions, Western blot analyses were performed for further quantification. In these analyses, we could confirm the previous observation that inflammatory conditions lead to increased integrin α11β1 levels, observable both in human and murine FLS. Specifically, RA-FLS showed up to 3.5 times increased levels compared to OA-FLS (n=5, two-tailed Mann–Whitney U test, n.s.). In hTNFtg FLS up to 13.8 times higher expression levels were detectable as compared to wt FLS (n= 4, two-tailed Mann–Whitney U test, p< 0.05) (Figure 1c). Next, we were interested to see if interactions between FLS and articular cartilage would affect the localization of integrin α11β1 and whether there are differences between wt and hTNFtg FLS. In an established *in vitr*o attachment assay [4], cocultures of hip caps from wt animals and FLS obtained from wt and hTNFtg animals were performed and analyzed by electron microscopy. Immunogold labelled particles detecting the anti-integrin α11β1 antibody demonstrated that in FLS-cartilage cocultures there were striking differences between the genotypes not only in the integrin α11β1 expression levels, but also in the localization of integrin α11β1. In hTNFtg FLS, a higher number of particles were detectable and most strikingly these were found particularly in areas with direct contact to the cartilage ECM in characteristic invading zones. Interestingly, some few particles were also found directly in the ECM itself. As an explanation for this phenomenon there might be also exist further FLS invasion zones which were not captured in these images. In contrast, very few particles were found in wt FLS with no prominent localization co cartilage contact areas (Figure 1d).

### *Itga11*^*-/-*^hTNFtg mice display an alleviated arthritic phenotype in comparison to hTNFtg mice

*Itga11*^*-/-*^hTNFtg mice were obtained by crossbreeding of *itga11*^*-/-*^ and hTNFtg mice. Mice were scored on a weekly basis by two independent observers assessing paw swelling and loss of grip strength. Both *itga11*^*-/*-^hTNFtg and hTNFtg mice displayed an onset of symptoms at around 5 weeks of age with increasing disease severity over time, as previously described [18]. *Itga11*^*-/-*^hTNFtg mice displayed a slightly alleviated phenotype with attenuated loss of grip strength and reduced paw swelling (Figure 2a). Wild type and *itga11*^*-/-*^ mice displayed no symptoms of inflammatory arthritis.

**Figure 2.**
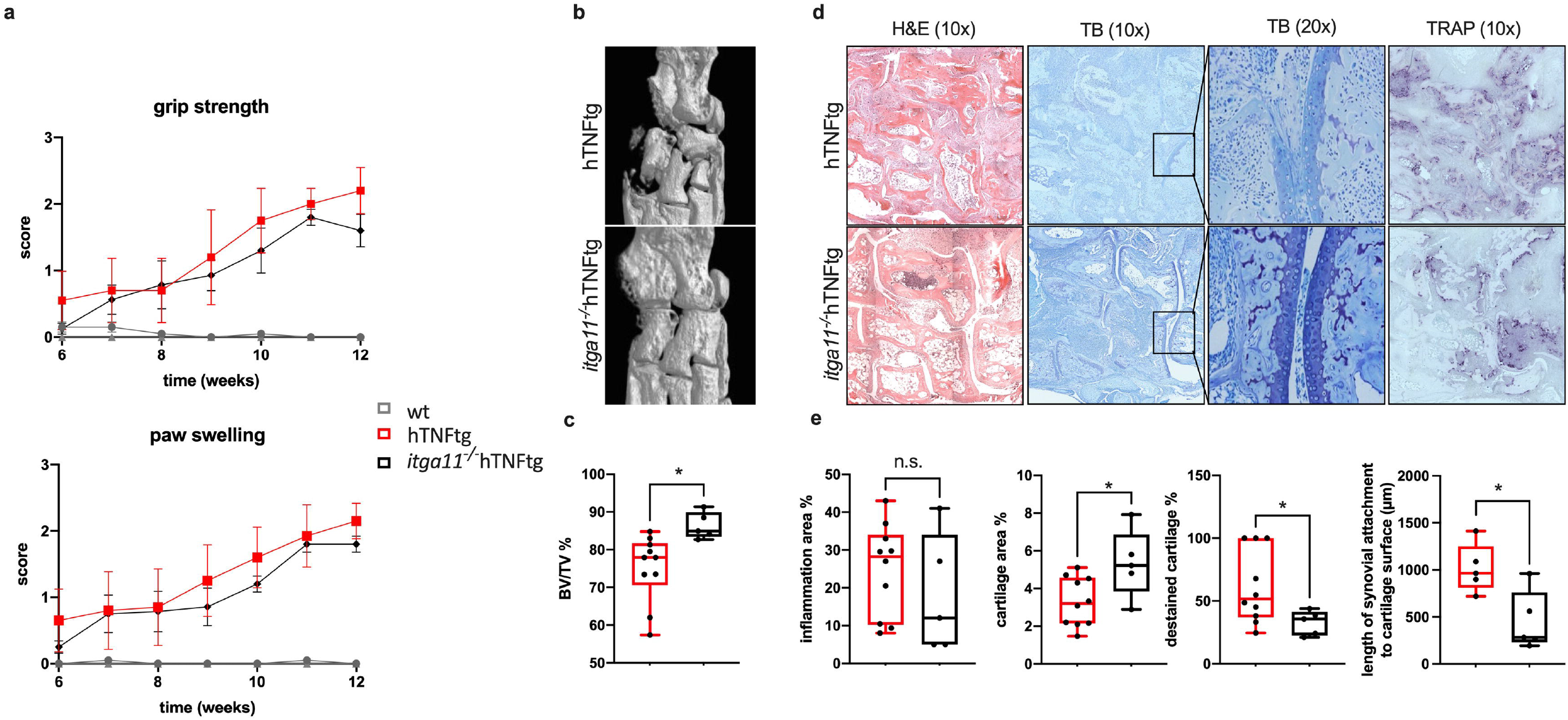
Effects of *itga11* knockout in hTNFtg mice. **a)** Clinical scoring of grip strength and paw swelling as surrogate parameters for inflammatory arthritis, as previously described [6] were assessed on a weekly basis (n ≥ 5). *Itga11*^*-/-*^hTNFtg mice displayed reduced severity of symptoms in comparison to hTNFtg mice from weeks nine to twelve. **b)** µCT imaging of hind paws showed less bone destruction in *itga11*^*-/*-^hTNFtg in comparison to hTNFtg mice at 12 weeks of age. **c)** Quantification of µCT imaging revealed higher residual bone volume of the second and third tarsal bone at twelve weeks of age in *itga11*^*-/-*^hTNFtg mice (+6.98% vs. hTNFtg, p < 0.005, two-tailed Mann–Whitney U test, n ≥ 5). **d)** H&E, Toluidine-blue and TRAP stainings were performed in decalcified, paraffin-embedded hind paws to visualize pathomorpholoical changes. *Itga11*^*-/*-^hTNFtg mice displayed visibly less joint destruction compared to hTNFtg mice. **e)** Histomorphological quantifications of Toluidine-blue stainings were performed in a blinded manner (n ≥ 5 animals per genotype). *Itga11*^*-/-*^hTNFtg mice displayed significantly more cartilage area (+2.02 % vs. hTNFtg, p < 0.05, two-tailed Mann–Whitney U test, n ≥ 5), less destained cartilage area (−16.08 % vs. hTNFtg, p < 0.05, two-tailed Mann–Whitney U test, n ≥ 5) and reduction in length of synovial attachment to cartilage surface (−806 µm vs. hTNFtg, p < 0.05, two-tailed Mann–Whitney U test, n ≥ 5), as markers for cartilage destruction. No significant differences between *Itga11*^*-/-*^hTNFtg and hTNFtg were found in the inflammation area (vs. hTNFtg, p > 0.05, two-tailed Mann–Whitney U test, n ≥ 5).

At the age of twelve weeks mice were sacrificed and µCT analyses of the hind paws were performed and analyzed qualitatively and quantitatively. In detail, less bone erosion was observed in hind paws from *itga11*^*-/-*^hTNFtg in comparison to hTNFtg mice (Figure 2b). Quantification of residual bone volume revealed significantly more bone volume in *itga11*^*-/-*^hTNFtg mice (+6.98% vs. hTNFtg, p < 0.005, two-tailed Mann–Whitney U test, Figure 2c), indicating less bone erosion in *itga11*^*-/-*^hTNFtg mice.

Histomorphological evaluations of joint pathologies were performed in H&E, Toluidineblue and TRAP stainings of paraffin-embedded sections and evaluated in a blinded manner (Figure 2d). Overall, *itga11*^*-/-*^hTNFtg mice showed less joint destruction, marked by more intact cartilage and less osteoclast formation in comparison to hTNFtg mice. In histomorphometric quantification of Toluidine-blue stainings *itga11*^*-/-*^hTNFtg mice showed a higher amount of total cartilage area (+2.02 % vs. hTNFtg, p < 0.05, two-tailed Mann–Whitney U test, Figure 2e) and a less destained cartilage (−16.08 % vs. hTNFtg, p < 0.05, two-tailed Mann–Whitney U test, Figure 2e) as markers for cartilage destruction. No significant differences between *itga11*^*-/-*^hTNFtg and hTNFtg were found in the inflammation area (vs. hTNFtg, p > 0.05, two-tailed Mann–Whitney U test). However, a significant reduction in length of synovial attachment to cartilage surface was found in *itga11*^*-/-*^hTNFtg mice (−806 µm vs. hTNFtg, p < 0.05, two-tailed Mann–Whitney U test, Figure 2e).

## Discussion

Integrin α11β1 is known to play an active role in CAFs in various tumor entities, in which it was associated with migration on, and remodeling of collagen [21]. With RA-FLS displaying a “tumor-like” behavior, our study establishes a link between integrin α11β1 and cartilage and bone destruction in RA in both human patients and an arthritis mouse model. So far integrin α11β1 has been described on cells of mesenchymal origin, including different types of fibroblasts [13, 19]. Several studies could also show that the expression of integrin α11β1 was upregulated by TGFβ and type 1 interferons, which are known to be involved in several autoimmune diseases and in RA-FLS transformation into the characteristic aggressive phenotype triggering further joint inflammation and destruction [22-25].

However, not all aspects of the functional role of integrin α11β1 in the context of RA have been understood so far. Although there is evidence that collagen citrullination negatively influences the adhesion of FLS by specifically decreasing the binding of integrin α11β1 to arginine-containing motifs thereby possibly modifying intracellular signaling in the pathogenesis of RA [26], analyses of the effects of integrin α11β1 deficiency on joint destruction and disease course have not been performed before. However, several studies were able to show the role of other integrins in RA-FLS associated with increased matrix binding, migration, proliferation, and cartilage destruction [5, 6, 28].

We found that integrin α11β1 was upregulated in both human RA patients as well as in hTNFtg mice in comparison to non-inflammatory controls such as OA patients and wild type mice. In human RA samples, the expression of integrin α11β1 was mainly located at the synovial sublining layer as shown by immunohistochemistry.

Interestingly, this was also observed in mice, but in addition, distinct staining clusters of integrin α11β1 were found in areas of pannus tissue adjacent to cartilage and bone at joint destruction sites. Immunofluorescence studies and Western Blot analyses confirmed the elevated expression of integrin α11β1 in human and murine FLS under arthritic conditions indicating an inflammation-induced upregulation of integrin α11β1 in FLS. Supporting a role for integrin α11β1 in FLS mediated cartilage destruction, the subcellular expression pattern showed integrin α11β1 localization primarily at sites of focal adhesion and cellular invasion. This was also demonstrated in our *in vitro* co-culture studies in which hTNFtg FLS were seeded onto cartilage explants and analyzed by transmission electron microscopy.

These results are in accordance with some published literature on the functional effects of integrin α11β1 outside the context of RA. In analogy to other collagen-binding integrins, integrin α11β1 was found to mediate fibroblast adhesion, cell migration and collagen reorganization as well as contraction and cell survival on collagen matrices leading to reduced proliferation and reduced adhesion to collagen in its absence [10, 29, 30]. In summary, the available literature suggests that integrin α11β1 mediates cell survival, adhesion to matrix, migration, and matrix-remodeling in fibroblasts, which are all essential characteristics for joint destruction mediated by RA-FLS in RA.

In our *in vivo* studies, knockout of *itga11* in the hTNFtg background resulted in the alleviation of the inflammatory arthritic phenotype as shown by clinical scoring, µCT imaging, and histological staining. Less cartilage destruction was observed in histomorphometric analyses of *itga11*^*-/-*^hTNFtg mice in comparison to hTNFtg animals. In detail, higher residual cartilage area after twelve weeks, and less destaining of articular cartilage, a surrogate for proteoglycan loss, were found in *itga11*^*-/-*^hTNFtg mice. Additionally, significantly less synovial attachment to cartilage was found in these mice. Attachment of RA-FLS to cartilage is a long-known hallmark feature in the induction of invasive cartilage destruction [4, 31]. Similarly, absence of the collagen-binding integrin α2β1 in antigen-induced arthritis (AIA) and hTNFtg mice, was shown to reduce the attachment of FLS to cartilage and cartilage destruction overall [6].

Furthermore, *itga11*^*-/-*^hTNFtg mice displayed elevated residual bone volume at twelve weeks of age, as well as reduced osteoclast formation in TRAP stainings compared to hTNFtg animals, indicating that less bone degradation takes place in this genotype. Although FLS can trigger bone erosions indirectly [32], they are not the primary cell line responsible for bone degradation. However, RA-FLS were shown to strongly promote osteoclastogenesis from precursor cells by the RANKL-RANK pathway [33, 34], suggesting an indirect effect that will require further investigations. As a further potential explanation, integrin α11β1 has been suggested to be involved in maintaining adult skeletal bone mass via binding to Osteolectin/Clec11a on skeletal stem cells and other osteogenic progenitors in bone marrow. This notion has been derived from data showing that deletion of *itga11* from bone marrow stromal cells impaired osteogenic differentiation and reduced osteogenesis and accelerated bone loss during adulthood in mice and men [35] which will be of interest for further studies.

Integrins were previously proposed as a viable target for the treatment of RA [36,37]. This study shows, that integrin α11β1 plays a role in RA, and that absence of the molecule leads to partial reduction of hallmark features of RA. Further studies into the exact molecular mechanisms of how integrin α11β1 contributes to the aggressive phenotype of RA-FLS will be needed, to assess the potential of integrin α11β1 as a therapeutic target.

## Conclusion

The results of this study suggest that the collagen-binding integrin α11β1 is upregulated in the context of RA and plays a role in adhesion and migration of fibroblasts on collagen and triggers cartilage degradation and destruction. The absence of integrin α11β1 in the hTNFtg mouse model leads to an alleviated phenotype, marked by reduced bone erosion and cartilage destruction, making the molecule a potentially interesting target for future therapeutical intervention studies.

## Acknowledgements

This work was supported by grants of the German Research Council to BE, TP, AKP (FOR2722) and BE (RU 2722, project ID 407239409). We thank George Kollias for providing us hTNFtg mice.

