## Supplement Table 1 for "Collagen-binding integrin α11β1 contributes to joint destruction in arthritic hTNFtg mice"

**Table 1** Primers used for genotyping

| **Primer name** | **Sequence (5' to 3')** | **Melting temperature (°C)** |
| --- | --- | --- |
| ITGA11 LacZ1 | GTGGTGGTTATGCCGATCGC | 63 |
| ITGA11 LacZ2 | TACCACAGCGGATGGTTCGG | 63 |
| ITGA11 ko-wt/F | CCATCAGAAGACAGGAGACGTATACAA | 67 |
| ITGA11 ko-wt/R | TGGTCAGTGGATGGGTTAGGAAG | 65 |
| TNF transgen fw | TACCCCCTCCTTCAGACACC | 62 |
| TNF transgen rev | GCCCTTCATAATATCCCCCA | 62 |
| TNF wt fw | GAGGGCCGAAGCTGCGGCTGGGT | 68 |
| TNF wt rev | GGTGGCGATTGGCTTGGCGAAG | 68 |
